# The Arabidopsis LHT1 amino acid transporter contributes to *Pseudomonas simiae*-mediated plant-growth promotion by modulating bacterial metabolism in the rhizosphere

**DOI:** 10.1101/2022.12.16.520833

**Authors:** Israel D. K. Agorsor, Brian T. Kagel, Cristian H. Danna

## Abstract

The root microbiome structure ensures optimal plant-host health and fitness, and it is, at least in part, defined by the plant genotype. It is well-documented that root-secreted amino acids promote microbial chemotaxis and growth in the rhizosphere. However, whether the plant-mediated re-uptake of amino acids contributes to maintaining optimal levels of amino acids in the root exudates, and in turn, microbial growth and metabolism, remains to be established. Here we show that LHT1, an amino acid inward transporter expressed in *Arabidopsis thaliana* roots, limits the growth of the plant-growth-promoting bacteria *Pseudomonas simiae* WCS417r (*Ps* WCS417r). Amino acid profiling of the *lht1* mutant root exudates showed increased levels of glutamine, among other amino acids. Interestingly, *lht1* exudates or Gln-supplemented wild-type exudates enhance *Ps* WCS417r growth. However, despite promoting bacterial growth and robust root colonization, *lht1* exudates and Gln-supplemented wild-type exudates inhibited plant growth in a *Ps* WCS417r-dependent manner. Transcriptional analysis of defense and growth marker genes revealed that plant growth inhibition was not linked to the elicitation of plant defense, but likely to the impact of *Ps* WCS417r amino acids metabolism on auxin signaling. These data suggest that an excess of amino acids in the rhizosphere impacts *Ps* WCS417r metabolism which in turn inhibits plant growth. Together, these results unveil that LHT1 regulates the amino acid-mediated interaction between plants and *Ps* WCS417r and suggest a complex relationship between root-exuded amino acids, root colonization by beneficial bacteria, bacterial metabolism, and plant growth promotion.

## INTRODUCTION

Crop agriculture has been fueled by an over-reliance on chemical fertilizers that have environmentally damaging consequences. Proposed alternative approaches to enhancing plant growth include the use of plant growth-promoting bacteria (PGPB) such as Pseudomonas simiae WCS417r, formerly known as *Pseudomonas fluorescens* WCS417r (1). In addition to factors that impact soil quality, metabolites exuded by roots have a major impact on the establishment of root associations with PGPB. Plants exude nutrient-rich fluids through their roots. These fluids, known as root exudates, influence the rhizosphere by inhibiting the growth of harmful microbes while promoting the growth of beneficial ones. Amino acids (AAs) are among the most represented metabolites in the root exudates of the model plant *Arabidopsis thaliana* (2–4). Microbes in the rhizosphere use AAs, or their derivatives, as food sources and perhaps to communicate with other microbes (*e.g.,* quorum sensing) (5–7) In addition, microbes use AA concentration gradients to navigate the soil toward roots. For example, in the colonization of tomato root tips, wild-type *Pseudomonas fluorescens* Pf0-1 outcompetes a triple-mutant derivative strain with impaired chemotaxis towards AAs (8). Microbes also use rhizospheric AAs to communicate with the plant. For instance, some microbial products trigger the release of AAs from plant roots (9), creating a self-reinforcing microbe-plant interaction. Microbes can also convert AAs into bioactive compounds with opposing effects on plant physiology. For example, microbes can convert AAs into plant growth-promoting substances such as the auxin hormone indole-3-acetic acid (IAA) (10). In contrast, on the other side of the spectrum, microbes have the capacity to metabolize AAs present in the rhizosphere (*e.g.*, glutamine) into derivatives that can inhibit plant growth (*e.g.*, ammonium) (11, 12).

On the plant side, the uptake of inorganic and organic nitrogen (including AAs) through the plant roots has been thoroughly studied, and AA-importers mediating this uptake have been reported (13–17). There are at least 61 transporters with confirmed or putative AA uptake activity encoded in the genome of *Arabidopsis thaliana* (18–20). However, to our knowledge, only three of these transporters localize to the plasma membrane of root cortex cells and can take up AAs at the low concentrations at which they are present in the soil. These three AA-importers are LHT1 of the LHT family (Lysine-Histidine-like Transporters), AAP5 of the AAP family (Amino Acid Permeases), and ProT2 of the ProT family (Proline Transporters) (21–25). These AA-importers have broad and overlapping specificities. However, they also show increased affinity for AAs that share molecular geometries and charges (21, 26, 27). Notwithstanding their detailed characterization, it is still unclear to which extent AA-importers expressed in the root cortex modulate the concentration of plant-derived AAs in the rhizosphere and how their activities contribute to building associations with beneficial bacteria. Answering this question is important as the concentration and balance of different AAs in the rhizosphere may impact the growth of belowground microbes, and hence, the growth of plants themselves. For example, colonization of cucumber roots by *Bacillus amyloliquefaciens* SQR9 enhances tryptophan secretion from the roots and promotes IAA biosynthesis by *B. a.* SQR9, which in turn boosts plant growth (10). Amino acids also serve as chemoattractants for different soil microbes such as *Pseudomonas fluorescens* Pf0-1 (8), *Ralstonia pseudosolanacearum* (28), and *Sinorhizobium meliloti* (29). To address if and how AA importers contribute to maintaining the amino-acidic composition of the root exudates, the composition and concentration of individual AAs were assessed in root exudates of Arabidopsis wild-type and loss-of-function mutants for the amino acid transporters AAP5, ProT2, and LHT1. Data obtained from AAs profiling, root colonization, plant growth assessment, and Arabidopsis gene expression indicate that LHT1 modulates the concentration of AAs in the root exudates, which in turn controls the growth and metabolism of *P.s. WCS417r* and its ability to promote plant growth.

## RESULTS

### The amino acid transporter LHT1 modulates amino acid content in Arabidopsis root exudates

Plants can modulate amino acid (AA) content in the rhizosphere by controlling the expression and activity of exporters that exude AAs out of the roots or importers that take up AAs present in the soil (30). However, it is less clear to which extent plants use AA-importers to take up AAs that the plants themselves exude into the rhizosphere. This putative retrieval of AAs from the exudates could contribute to modulating AA content in the rhizosphere, and hence, the growth of PGPB. To establish whether plants control the accumulation of plant-derived amino acids in the rhizosphere through the activity of AA importers, loss-of-function mutants for AA transporters with confirmed uptake activity and expressed in the root (21–25) were tested. The loss-of-function mutants *aap5, prot2,* and *lht1* were chosen because: (i) they represent three families of AA importers, (ii) are expressed in the cortex of the root (23, 25, 27), and (iii) LHT1 and AAP5, at least, can take up AAs at the concentrations present in the soil (24).

The experimental strategy was based on the method described by Haney *et al* (31) with some modifications (**Fig. S1A**). Briefly, Arabidopsis seedlings were initially grown in full-strength (1x) MS medium with sucrose (0.5%) for 12 days and then transferred to half-strength (0.5x) MS without sucrose for 3 days. To enable the collection of root exudates for further analysis, the seedlings were grown with the roots submerged in liquid culture and separated from the shoots by a sterilized polytetrafluoroethylene (PTFE) mesh that floats at the surface of the liquid. After 15 days of growth, the root exudates were collected, and the total AA content was assessed using as previously described (32). Two *aap5* and one *prot2* mutant lines showed no differences in total AA content in the root exudates when compared to Col-0 wild-type plants (**Fig. S1B**). By contrast, the root exudates of the *lht1* mutants showed increased levels of AAs when compared to wildtype (**Fig. 1A**). Since LHT1 localizes to the plasma membrane of the rhizodermis (21) where it acts as an AA importer (22) and the culture media was not supplemented with AAs, the AAs that accumulate in the *lht1* root exudates must be of plant origin. Hence, the changes in the composition of the *lht1* exudates are most likely due to deficient rhizospheric AA reuptake by LHT1. Together, the results indicate that the loss of the AA importer LHT1 can alter the AA profile of root exudates and that not all AA importers expressed in the root serve the function of maintaining the AA concentrations of the exudates. Even though the biochemical activities and localization of LHT1 have been known to be compatible with regulating the rhizosphere’s AA composition through re-uptake, this specific function had not been previously reported.

**Figure 1.**
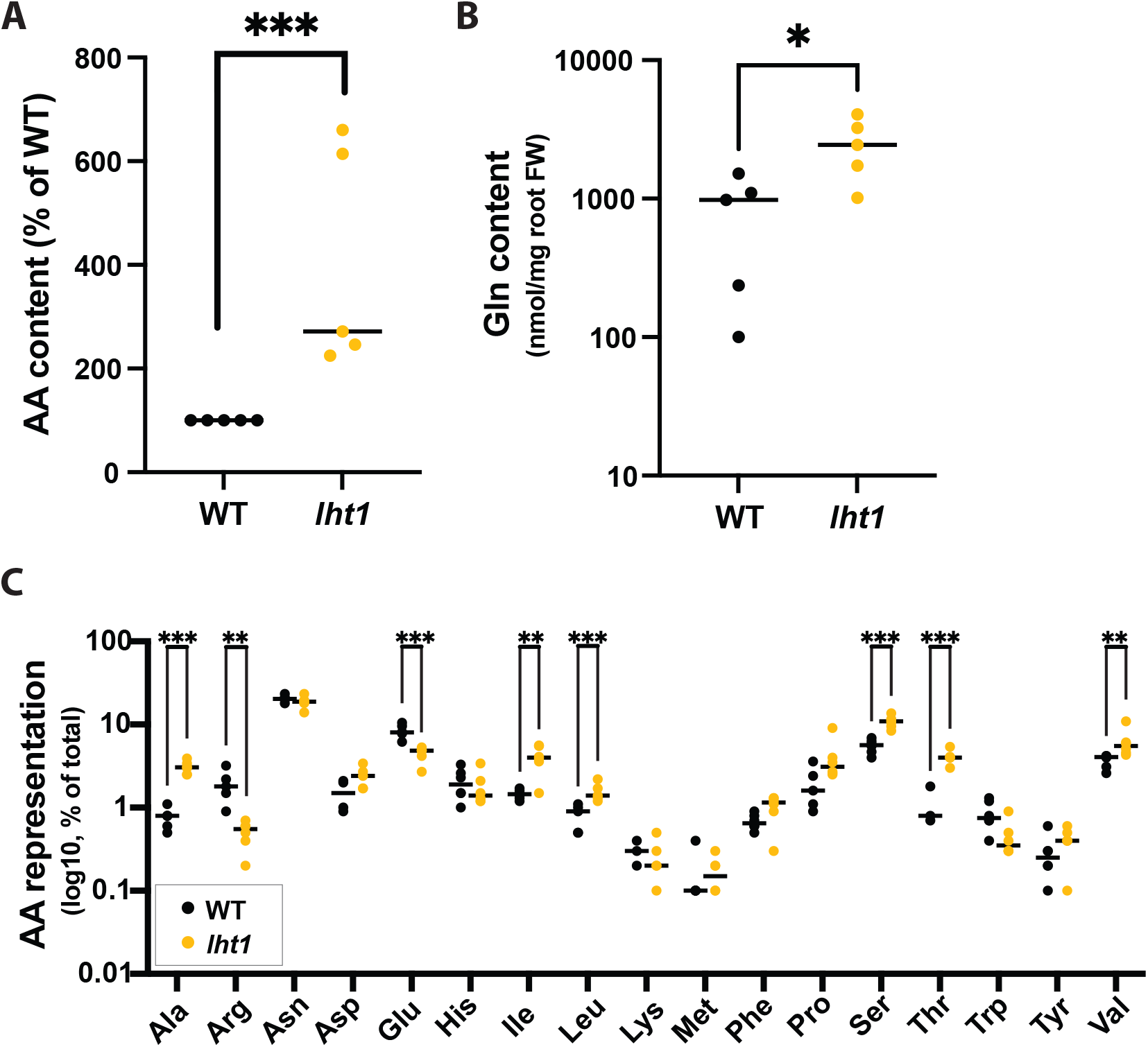
LHT1 modulates amino acid content of Arabidopsis root exudates. Across all panels in this figure, unless otherwise stated, data are from at least 3 independent biological replicates. In each biological replicate, at least 5 seedlings/plants or their products (e.g., exudates) were analyzed. Each dot in the scattered dot plots shows the median of the 5 or more seedlings/plants, and the median of all replicates is depicted as a bar. A two-sided Student’s t-test was performed for statistical comparison of two means, or a Welch’s t-test for two means with unequal variances, when relevant. P values are represented as *≤0.05, **≤0.01, and ***≤0.005. Further experimental and analysis details are described in the Methods section. (A) lht1-mutant root exudates have higher AA content than wild-type (WT) exudates. Exudates obtained from 15-day-old lht1 or WT seedlings grown on 1x MS liquid medium containing 0.5% sucrose were filter-sterilized, and their total AA content was assessed using the L-Amino Acid Quantitation Colorimetric/Fluorometric Kit (see Methods). Each data point represents the median of 6 wells in each of 5 independent biological replicates. Data originally scored as nM of total L-AAs per mg of fresh root tissue are presented as a percentage of WT to be able to compare independent biological replicates in the same graph. (B) lht1-mutant root exudates have higher glutamine (Gln) content than wild-type exudates. Root exudates obtained as in (A) were treated with formic acid and profiled via LC-MS. Data analysis was performed on 5 independent samples per condition. (C) lht1-mutant root exudates show imbalanced AA content. Data obtained as in (B) is shown as a percentage for total AAs in lht1 and WT exudates. Glycine data was omitted because unplanted MS medium contained background levels of glycine (2 mg/L). LC-MS analysis was performed on 6 independent samples per condition.

Liquid chromatography coupled to mass spectrometry (LC-MS) analysis was used to define the specific AA changes in the root exudates of the *lht1* plant. This analysis revealed that glutamine (Gln) was the most abundant amino acid that had significantly elevated concentrations in the root exudates of *lht1* compared to those of wild-type plants (**Fig. 1B**). Nonetheless, in line with the broad AA-substrate specificity of LHT1 (21), the analysis revealed that several other AAs were enriched in *lht1* root exudates, including alanine, asparagine, aspartic acid, leucine, isoleucine, methionine, phenylalanine, proline, serine, threonine, and valine when compared to the root exudates of wild-type plants (**Fig. 1C**). The rationale to assign LHT1 a role in AA re-uptake from the root exudates is based on its thoroughly established importer function (33). Since LHT1 moves AAs only in an inward direction and our root exudate assays were performed on AA-free media, the most parsimonious model to explain the overaccumulation of AAs in the *lht1*-root exudates is that LHT1 modulates the levels of AAs in the exudates through the re-uptake of previously exuded amino acids.

### *lht1* root exudates enhance *Ps* WCS417r growth

Plant-Growth-Promoting Bacteria (PGPB), such as *Ps* WCS417r (formerly *Pseudomonas fluorescens* WCS417r), colonize the surface of Arabidopsis roots and promote plant growth (1). Since the growth of PGPB depends on the biophysical properties and composition of the rhizosphere, which are modulated by the plant itself, testing whether the changes in *lht1* root exudates affected the growth of *Ps* WCS417r would provide important information to define which plant-made AAs could favor bacterial and plant growth. To this end, the growth of *Ps* WCS417r in wild-type and *lht1* root exudates, as well as in unplanted MS media (MS medium not conditioned by plants) as a control, was longitudinally measured using absorbance OD_600nm_ over a 24 h period with intermittent shaking in a microtiter-plate reader. In line with their higher AA content, *lht1* root exudates supported more bacterial growth than wild-type root exudates (**Fig. 2A**). Considering that wild-type and *lht1* root exudates may differ in the content and abundance of metabolites other than AAs, testing the effect of AA abundance on PGPB growth would at least address if AAs that are over-represented in *lht1* exudates could augment bacterial growth. Supplementation of wild-type exudates with Gln was sufficient to boost *Ps* WCS417r growth (**Fig. 2B**), suggesting that plant-derived Gln is a limiting factor for *Ps* WCS417r growth in the Arabidopsis rhizosphere. Similarly, supplementing wild-type root exudates with serine (Ser), another AA that accumulates in the *lht1* root exudates, promoted *Ps*WCS417r growth (**Fig. 2C**), suggesting an overall positive correlation between amino-acid content in the root exudates and the growth of beneficial microbes in the rhizosphere.

**Figure 2.**
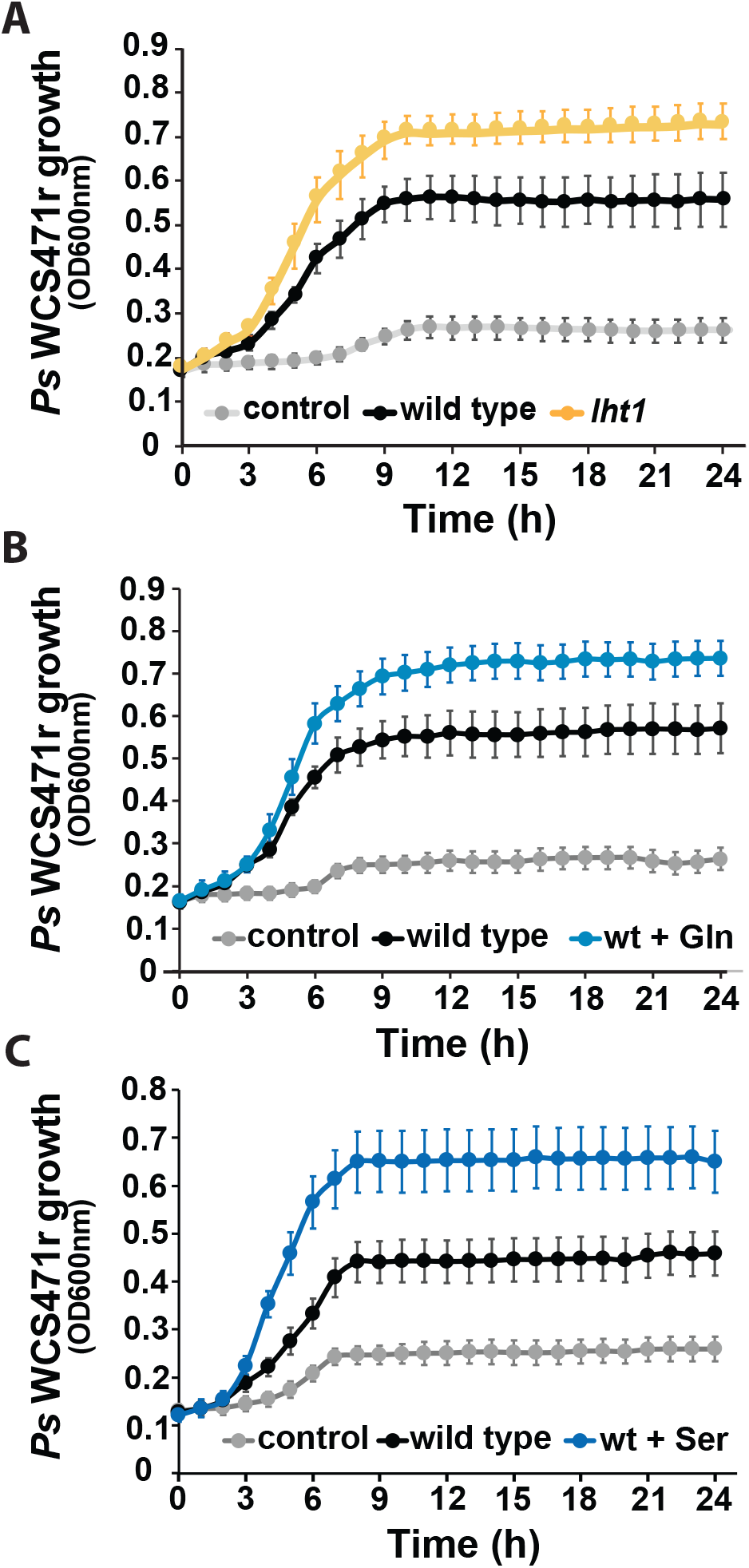
lht1-derived root exudates enhance Ps WCS417r growth. Across all panels in this figure, data represent 3 independent biological replicates. Exudates were obtained as described in Fig. 1A and Methods. In each biological replicate, at least 6 wells per condition were seeded with exudates or control media and longitudinally scored for growth every hour using OD600nm as the readout. Plates were incubated and scored in a temperature-controlled plate reader with shaking. Control corresponds to MS medium without exudate. Each dot in the curves represents the median growth of 6 wells at a given time point. Error bars are SEM. Statistical analysis was done using the CGGC (Comparison of Groups of Growth Curves) permutation test to compare pairs of samples (e.g., wild type vs. lht1) over the course of growth (24 hours). All differences between conditions were statistically significant. Further experimental and analysis details are described in the Methods section. (A) Ps WCS417r grows better in lht1-derived than WT exudates. (B) Ps WCS417r grows better in Gln-supplemented WT exudates. WT exudates were supplemented with 1 mM Gln (final concentration) or an equal volume of water as control. (C) Ps WCS417r grows better in Ser-supplemented WT exudates. WT exudates were supplemented with 1 mM Ser (final concentration) or an equal volume of water as control.

### Loss of *lht1* promotes root colonization by *Ps* WCS417r

Having uncovered that *lht1* root exudates promote more *Ps* WCS417r growth, the next question to address was whether this enhanced *in vitro* growth would translate to better root colonization. To this end, Arabidopsis seedlings were grown for two weeks in MS-PhytoAgar plates incubated vertically and then transferred to sterile plates with autoclaved-3MM paper as the substrate (**Fig. S2A & B**). One day later, roots were flood-inoculated with *Ps* WCS417r. After inoculation, the plates were incubated horizontally, and root colonization proceeded for 72h. At that time, roots were weighed and processed to assess colonization as colony-forming units (CFUs) per root weight. In line with the *in vitro* growth assays, the roots of *lht1* supported more bacterial growth than the roots of wild-type seedlings (**Fig. 3A**). These data support the hypothesis that the higher concentrations of AAs present in the rhizosphere of *lht1* plants have a positive impact on the growth of PGPB such as *Ps* WCS417r. Next, to investigate the mechanisms underlying the enhanced bacterial colonization of *lht1* roots, two processes known to contribute to the establishment of a successful root-bacteria interaction, chemotaxis (8, 34, 35) and biofilm formation (36, 37), were tested. The chemotaxis index was assessed using a modified capillary assay previously described (34, 38) (**Fig. S3**). Unplanted MS media (MS medium that has not been conditioned by roots) was used to assess the background swarming motility of *Ps* WCS417r. The data showed a significant preference for *lht1* root exudates in both non-competitive (**Fig. 3B**) and competitive chemotaxis assays (**Fig. 3C-E**). To define whether the chemotaxis-enhancing capacity of *lht1* exudates was driven, at least in part, by the AAs that promote bacterial growth, wild-type exudates supplemented with Gln were also tested. Indeed, wild-type exudates supplemented with Gln showed higher chemotaxis capacity and hence higher *Ps* WCS417r titers than non-supplemented wild-type exudates (**Fig. 3F**). The capacity of *Ps* WCS417r to form biofilms in wild-type and *lht1* root exudates, as well as in unplanted media as a control, was assessed using the crystal violet staining method (37). We hypothesized that the increased accumulation of AAs observed in the root exudates obtained from *lht1* plants could facilitate biofilm formation and, in this way, the growth of *Ps* WCS417r in the rhizosphere. Consistent with this hypothesis, *Ps* WCS417r showed an enhanced capacity to form biofilm in the *lht1* root exudates compared to wild-type root exudates (**Fig. 3G**). Therefore, altogether, the AA transporter LHT1 defines the AA content of the root exudates, the chemotaxis, the biofilm formation, and the growth capacity of *Ps* WCS417r.

**Figure 3.**
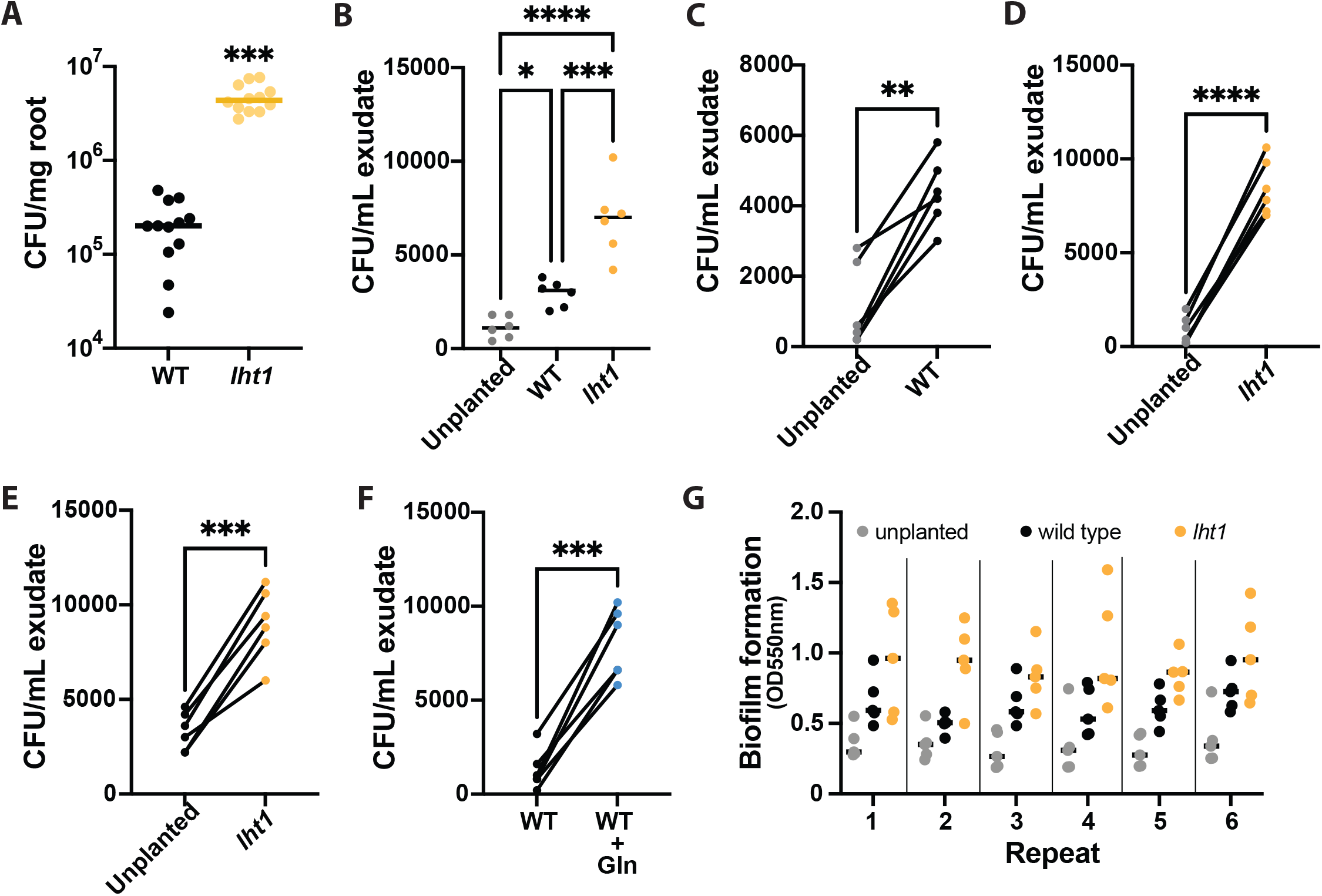
Ps WCS417r more effectively colonizes lht1 than wild-type roots. Across all panels in this figure, data are from at least 5-6 independent biological replicates. In each biological replicate, at least 5 seedlings/plants or their products (e.g., exudates) were analyzed. Each dot represents the median of 5 seedlings, 5 independently collected exudates, or 3 chemotaxis syringes per condition. In panels (A) the median of all biological replicates is shown as a bar. A two-sided Student’s t-test was performed for statistical comparison of two means or a Welch’s t-test for two means with unequal variances. For comparison of more than two means, a one-way ANOVA followed by Tukey’s posthoc test, or a Kruskal-Wallis test for unequal variances followed by Dunn’s posthoc test was performed. P values are represented as *≤0.05, **≤0.01, ***≤0.005, and ****≤0.001. Further experimental and analysis details are described in the Methods section. (A) Loss of LHT1 promotes root colonization by Ps WCS417r. The 1x MS-wetted 3MM paper in which 6 seedlings were growing was inoculated with Ps WCS417r at OD600nm= 0.00002. Seventy-two hours later, the roots were harvested, weighed, and ground; serial dilution were plated to count colony forming units (CFU) on LB agar with 50 μg/mL of rifampicin. Colonization is presented as CFU per mg of root fresh weight. (B) AAs-rich lht1 exudates attract Ps WCS417r more effectively than WT exudates. For this non-competitive chemotaxis assay, 200μl of filter-sterilized exudates of each condition were loaded in 1 mL syringes without the needles. The tip of each syringe was then dipped in an independent Petri dish containing chemotaxis buffer inoculated with Ps WCS417r at a final OD600nm= 0.002. After 1h, bacterial cells inside the syringe were harvested and diluted to measure CFU. Data are depicted as total CFU per mL of exudate. (C-D) Gln-supplemented WT exudates attract Ps WCS417r more effectively than non-supplemented WT exudates. For these competitive chemotaxis assays, 200 μL of filter-sterilized exudates of each condition were loaded in 1 mL syringes. Then, the tip of two syringes of different conditions was dipped in the same Petri dish containing chemotaxis buffer inoculated with Ps WCS417r at a final OD600nm= 0.002. After 30 min, bacterial cells inside each of the two syringes were harvested and diluted to measure CFU. Data are presented as total CFU per mL of each exudate. Competitive chemotaxis results are presented as follows: (C) Control 0.5x MS medium (without sucrose) versus WT exudate, (D) Control 0.5x MS medium (without sucrose) versus lht1 exudate, (E) WT versus lht1 exudate, and (F) WT exudate versus WT exudate supplemented with 1 mM Glutamine. (G) lht1 exudates better support Ps WCS417r capacity to produce biofilm. Six aliquots of 0.5x MS medium (without sucrose), wild-type root exudates, or lht1 root exudates were loaded on independent wells of a 96-well round-bottom plate and then inoculated with 2 μl freshly harvested Ps WCS417r grown overnight in LB medium After 48h of static growth, biofilms were stained with crystal violet, and absorbance at 550 nm was measured in a microplate reader as described in Methods. Each depicted dot represents the absorbance in each of 6 wells per condition. Biofilm formation was independently tested 6 times.

### Rhizospheric amino acids stimulate robust root colonization but compromise plant growth in a microbial-dependent manner

*Ps* WCS417r colonizes the root surface and promotes the growth of wild-type plants with a wild-type rhizospheric composition. Since the AA-enriched exudates of *lht1* support more bacterial growth than wild-type exudates, we hypothesized that wild-type plants exposed to *lht1* exudates and *Ps* WCS417r (*lht1*+Ps) would grow larger than plants exposed to wild-type exudates and *Ps* WCS417r (WT+Ps). Unexpectedly, this analysis showed the opposite outcome: *lht1* exudates and *Ps* WCS417r inhibited the growth of wild-type seedlings (**Fig. 4A**). Importantly, as expected, when wild-type seedlings were treated with *Ps* WCS417r only, they grew larger than untreated seedlings. Similarly, *lht1* exudates alone did not inhibit the growth of wild-type seedlings (**Fig. 4A**). Therefore, the growth inhibition depends on both *lht1* root exudates and *Ps* WCS417r. To narrow down the mediators of this plant growth inhibitory effect, Arabidopsis growth was assessed after exposure to Gln-supplemented wild-type root exudates and *Ps* WCS417r (WT-Gln+Ps). Like *lht1+Ps,* WT-Gln+Ps also inhibited the growth of wild-type Arabidopsis seedlings (**Fig. 4B**). Ruling out a direct toxic effect of Gln, seedling growth was not inhibited by 1 mM or 10 mM Gln (**Fig. 4B**), implying a microbe-mediated inhibition of plant growth that is dependent on *Ps* WCS417r Gln metabolism. Importantly, a high *Ps* WCS417r inoculation titer (OD_600nm_ =1.6) did not compromise plant growth, suggesting that the high bacterial colonization in *lht1+Ps* and WT-Gln+Ps does not contribute to plant growth inhibition (**Fig 4B**). To test the impact of other AAs that accumulate in *lht1* root exudates, plant growth in wild-type exudates supplemented with 1 mM or 10 mM serine (Ser) was assessed. Interestingly, an excess of Ser significantly inhibited plant growth at both concentrations tested and in the absence of *Ps* WCS417r (**Fig. 4C**). These data demonstrate that Ser does not contribute to the plant growth inhibitory effect of *lht1* exudates mediated by *Ps* WCS417r.

**Figure 4.**
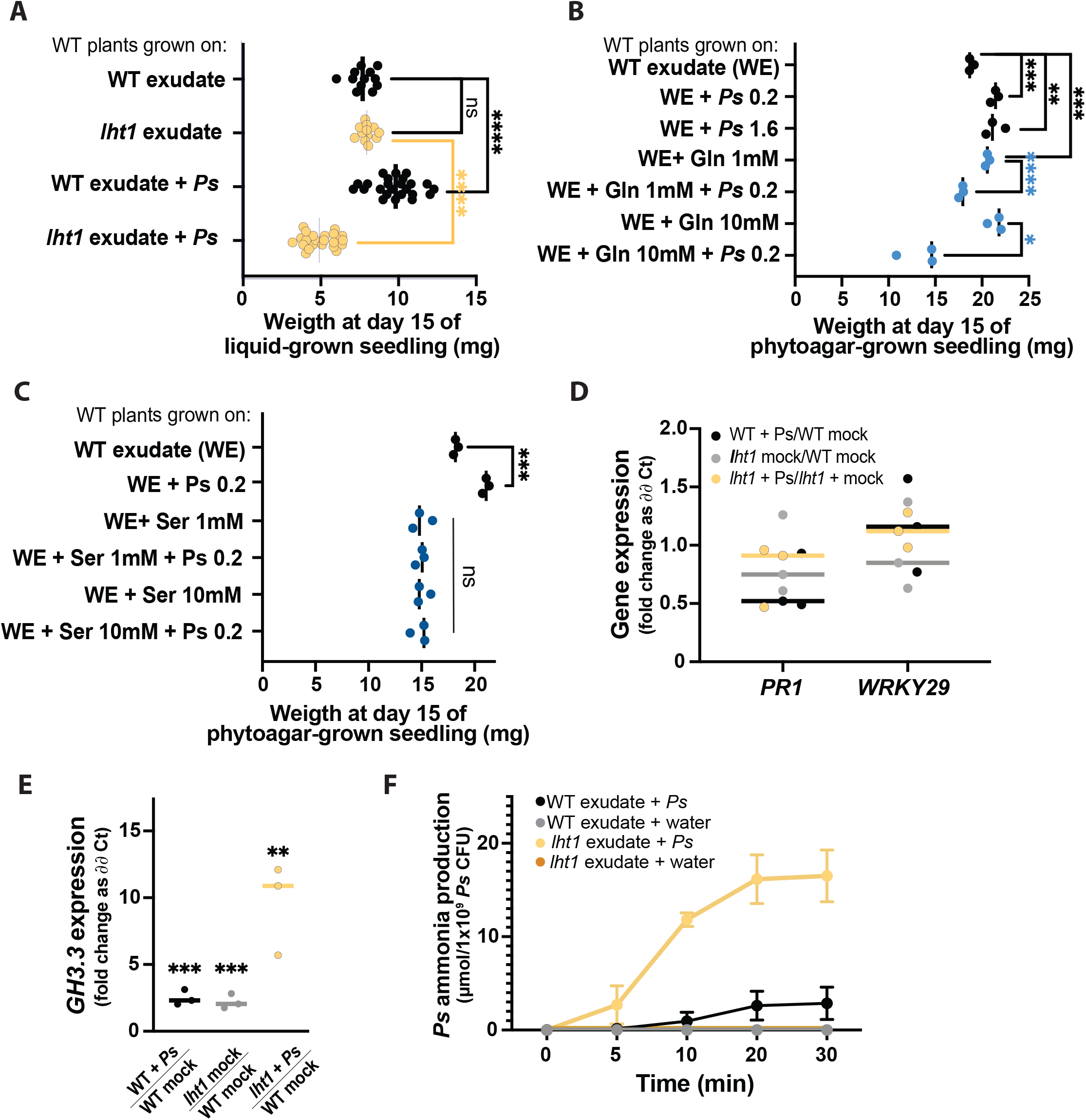
Excess AAs in lht1 exudates combined with Ps WCS417r inhibit wild-type Arabidopsis growth. Across all panels, each dot represents an independent biological replicate. The median of all biological replicates is depicted as a bar. A two-sided Student’s t-test was performed for statistical comparison of two means or a Welch’s t-test for two means with unequal variances. For comparison of more than two means, a one-way ANOVA followed by Tukey’s posthoc test, or a Kruskal-Wallis test for unequal variances followed by Dunn’s posthoc test was performed. P-values are represented as *≤0.05, **≤0.01, ***≤0.005, and ****≤0.001. Further experimental and analysis details are described in the Methods section. (A) lht1 exudates inhibit the growth of wild-type Arabidopsis when inoculated with Ps WCS417r. Fifteen-day-old wild-type seedlings growing on 12-well plates with floating mesh on WT or lht1 exudate with or without Ps WCS417r OD600nm= 0.2. Each dot represents the median weight of ≥3 seedlings per condition. The experiment was repeated 6 times; the median of all biological replicates is depicted as a bar. (B) Gln-supplemented WT exudates combined with Ps WCS417r inhibit the growth of wild-type Arabidopsis. Assay was carried out as in (A) except that two different Ps WCS417r inoculum titters were used (OD600nm=0.2 or 1.6) and that WT exudates were supplemented with Gln 1mM or 10 mM and inoculated or not with Ps WCS417r (OD600nm=0.2). (C) Ser-supplemented WT exudates directly inhibit the growth of wild-type Arabidopsis. Assay and controls as in (B), except that WT exudates were supplemented with 1 mM or 10 mM Ser (instead of Gln). (D) Pathogen-responsive genes are not induced in plants concomitantly exposed to Ps WCS417r and lht1 exudates. Wild-type seedlings were exposed for 24h to WT exudate (depicted as WT mock), WT exudate + Ps WCS417r (final OD600nm= 0.2), lht1 exudate (depicted as lht1 mock), or lht1 exudate + Ps WCS417r (final OD600nm= 0.2) as described herein (A) and in Methods. Gene expression in the roots was assessed using RT-qPCR. Fold change was calculated using ∂∂Ct. The specific comparisons are depicted in the X-axis. Each dot corresponds to an independent biological replicate (calculated based on 2 experimental replicates), and the median of the 3 biological replicates is depicted as a bar. (E) GH3.3, a mediator of the stress-triggered growth inhibition response, is induced in plants concomitantly exposed to Ps WCS417r and lht1 exudates. Expression of the gene encoding the auxin-conjugating enzyme GH3.3 is measured in the conditions described in (D). (F) Ps WCS417r produces more ammonia from lht1 than from wild-type exudates. Ammonia excreted by Ps WCS417r while incubated on WT or lht1 exudates was assessed at 0, 5, 10, 20, and 30 min. Exudates obtained in three independent experiments were assessed for ammonia independently but simultaneously. After the indicated times, the incubation-media supernatants were harvested, filter sterilized, and their ammonia content was assessed using an ammonia-assay kit (AbCam; Cat # ab102509) following manufacturer’s guidelines. Each dot represents the median ammonia concentration (μmol/mL), and error bars correspond to the SEM.

Across plant species, the response to abiotic and biotic stressors usually includes the inhibition of plant growth (39–41). To identify the underlying mechanisms associated with plant growth inhibition when *Ps* WCS417r colonizes an AA-rich rhizosphere, the expression of Arabidopsis genes induced in response to various stressors that produce plant growth inhibition, was tested. First, the elicitation of plant immunity is typically accompanied by the inhibition of plant growth (42–44). To test if plant immunity was elicited by the combination of *lht1* exudates and *Ps* WCS417r, the expression levels of *PR1* and *WRKY29* were assessed by RT-qPCR. The expression of these genes is induced in response to a broad range of pathogenic cues (45, 46). Arguing against a generalized activation of the plant defense in response to larger-than-normal root colonization that could negatively impact plant growth, the data showed similar levels of *PR1* or *WRKY29* expression in wild-type roots colonized with *Ps* WCS417r regardless of the origin of the plant exudates (**Fig. 4D**). The levels and potency of the plant growth hormone indole-3-acetic acid (IAA) are partially controlled by the activity of enzymes that conjugate amino acids to IAA, producing IAA-derivatives less capable of promoting plant growth (47). Some members of the GH3 family of IAA-amido synthetases whose encoding genes transcriptionally respond to stress are necessary and sufficient to promote stress-induced plant growth inhibition, presumably by lowering the effective concentration of active IAA (47). *GH3.3,* in particular, has been used as a marker of stress-mediated growth inhibition (48). The expression of *GH3.3* in wild-type plants exposed to *lht1* exudates and *Ps* WCS417r was significantly higher than that of wild-type plants exposed to WT exudates and *Ps* WCS417r (**Fig. 4E**). Lastly, as excessive ammonia is known to inhibit plant growth (11, 12, 49), we tested the hypothesis that *Ps* WCS417r may convert the excess Gln present in *lht1* exudates or Gln-supplemented WT exudates into ammonia. Thus, the capacity of *Ps* WCS417r to produce ammonia when grown in wild-type or *lht1* exudates was assessed. In line with the ammonia-toxicity hypothesis, the levels of ammonia produced by *Ps* WCS417r were higher in the *lht1* exudates than in the WT exudates (**Fig. 4F**). Notwithstanding the lack of direct evidence, the increased levels of AAs in the *lht1* root exudates could enhance *Ps* WCS417r-mediated production of ammonia, which could, in turn, contribute to inhibiting plant growth. Overall, the data presented in this section suggest that LHT1-mediated regulation of the root exudate’s amino-acidic composition is critical to maintaining the concentration and representation of AAs that is optimal for the beneficial effect of root colonization by PGPB.

## DISCUSSION

### LHT1 activity is essential to maintain a balanced composition of amino acids in the root exudates

Plants release a fraction of their metabolites into the rhizosphere as root exudates. Since some of the exudate-dependent microbes present in the rhizosphere promote plant growth, the metabolites present in root exudates represent an investment rather than an expenditure for the plant. It has been proposed that, like seeds or growing organs (e. g. new leaves), belowground microorganisms are additional sinks of plant photosynthates (50). Therefore, as with other plant sinks, it is reasonable to hypothesize that plants would regulate the transport of metabolites to and from the rhizosphere. Furthermore, it had been hypothesized that active recycling of low molecular weight compounds from the rhizosphere is particularly important for plants to control the growth of rhizosphere microbes and to ward off pathogen invasion of root tissues (51). Yet, whether plants exert control over the accumulation of plant-derived metabolites in the rhizosphere through AA uptake was unclear. The present study was based on two related hypotheses: (i) plants would regulate the levels of AAs in the rhizosphere, including the AAs released by the plants themselves, and (ii) some of the AA importers that contribute to the uptake of AAs from the growth medium and the soil, may also contribute to retrieve plant-derived AAs back into the plant. In support of these hypotheses, the data showed that a loss-of-function mutant of the AA importer LHT1 leads to an increased accumulation of AAs in the root exudates of *Arabidopsis* (**Fig. 1A, C**). Furthermore, our search for specific exuded metabolites, able to change the structure of the rhizosphere, identified Gln, among other AAs, enriched in the root-exudates of the *lht1* (**Fig. 1B**) whose sole supplementation is sufficient to boost the growth of the *Ps* WCS417r (**Fig. 2A, B**). Unexpectedly though, the increased bacterial growth in exudates and increased root colonization (**Fig. 3A**) did not translate into increased plant growth (**Fig 4A**). Together, the data suggest that wild-type plants maintain a balanced concentration of AAs in the exudates that is optimal to promote PGPB growth and, in turn, to maximize plant growth, and that LHT1 contributes to maintaining this optimal concentration of AAs in the exudates. In line with an optimized plant-microbe interaction that is, at least in part, modulated by LHT1, one hour after inoculation of Arabidopsis roots with *Ps* WCS417r, *LHT1* expression increases in root tissues (52), suggesting a mechanism by which microbes and plants communicate with each other to maintain levels of AAs in the rhizosphere that sustain optimal PGPB growth and metabolism, and enhanced plant growth.

### The re-uptake of root-exuded amino acids is necessary to modulate bacterial metabolism and promote plant growth

The root-associated microbiome is mainly located in the rhizosphere, the narrow (1-3 mm) region of soil surrounding the root that is rich in plant-made metabolites exuded by the roots (53). Several PGPB species that live in the rhizosphere have been shown to either produce plant hormones or regulate the levels of plant hormones that have a positive effect on plant growth (54, 55). The mechanisms by which *Ps* WCS417r promotes plant growth, however, are not fully understood. To explain the detrimental effects of Gln-supplemented *Ps* WCS417r on plant growth, several hypotheses can be formulated, some of which would be worth testing in future studies. One hypothesis would be that excessive levels of AAs could have a direct detrimental effect on the physiology of the plant itself. This may be proven correct for some AAs, including Ser, which was sufficient to inhibit Arabidopsis root and shoot growth (**Fig. 4C**). Gln has been reported to inhibit root growth of the Ler (Lansberg erecta) ecotype but not the Sha (Shakdara) ecotype of Arabidopsis (56), suggesting that the root growth of the Col-0 ecotype used in our experiments could also be inhibited by an excess of Gln. However, in the experimental setup used in our study, the detrimental effect of Gln was only evident in the presence of *Ps*WCS417r (**Fig. 4B**). For instance, even relatively high concentrations (20 mM) of Gln alone did not inhibit plant growth (**Fig. 4B)**. In contrast, the combination of *Ps*WCS417r with 1 mM Gln causes subtle growth inhibition, and 10 mM robustly inhibits plant growth (**Fig. 4B**). This observation suggests that the microbe-mediated bioconversion of Gln is required to inhibit plant growth. Bioconversion by soild-welling microbes has been extensively studied for chemicals of anthropogenic origin (*e.g.,* pesticides) (57, 58) but much less is known about PGPB bioconversion of plant-derived compounds. Nevertheless, it is well-documented that microbes convert Gln into ammonium, and that high ammonium levels inhibit the growth of roots and shoots in Arabidopsis (59), especially in conditions of low potassium levels (11, 12, 49). Furthermore, the genome of *P. simiae* encodes at least five enzymes capable of converting glutamine into glutamate and ammonia (www.Biocyc.org). This ammonia-producing reaction, together with the inhibition of the ammonia-consuming reaction carried out by glutamine-synthase, would be favored in the Gln-rich environment of the *lht1* rhizosphere. In agreement with the literature, *Ps* WCS417r produced high ammonia levels when *lht1* exudates were used as growth medium (**Fig. 4F**). However, in the conditions used for assessing plant growth in this study, the high availability of potassium in the MS medium may alleviate ammonia toxicity. Considering that the high ammonia level present in the MS medium (20 mM) does not inhibit plant growth, the local concentrations of ammonia produced by *Ps* WCS417r Gln metabolism would need to be at least higher than 20 mM to be able to inhibit plant growth. Thus, in future studies, it would be relevant to test whether *Ps* WCS417r strains with reduced capacity to release ammonium from glutamine (*e.g.,* a *glsA* mutant; locus tag PS417_15505) would maintain the capacity to grow robustly in exudates supplemented with Gln and perhaps promote plant growth to a larger extent compared to the wild-type strain. Other byproducts of PGPB metabolism may also affect plant growth; thus, in future studies, it would be informative to perform metabolic profiling of root exudates post-PGPB growth to identify metabolites that are only present at high levels in the supernatant of *Ps* WCS417r grown in *lht1* exudates. A complementary hypothesis would be that *Ps* WCS417r-generated byproducts of Gln metabolism activate a stress response in the plant that includes growth inhibition. Reduced plant growth in response to abiotic and biotic stress is well documented. The metabolism of plant growth hormones such as IAA mediates the growth inhibitory response to stress (47). The GH3 family of proteins are important player in the plant inhibitory metabolism of IAA upon stress. In line with the notion that the Gln metabolism by *Ps* WCS417r described here may promote the formation of IAA-derivatives that suppress plant growth, Gln-supplemented *Ps* WCS417r, but not Gln or bacteria alone, promoted the induction of the Arabidopsis gene encoding the IAA-amido synthetase GH3.3. This result provides a clue for future dissection of the mechanism by which Gln metabolism in *Ps* WCS417r inhibits plant growth.

## CONCLUSIONS

Data presented in this study establishes that Arabidopsis uses AAs reuptake to control the organic composition of the rhizosphere as a relevant strategy to foster enduring beneficial interactions with their root microbiota. Modulation of the concentration of AAs in the exudates is, at least in part, executed by LHT1. In the absence of LHT1-mediated re-uptake, the excessive levels of AAs in the rhizosphere cause microbe-mediated inhibition of plant growth. Future studies will define how Gln metabolism in *Ps* WCS417r leads to plant growth inhibition. Among other possibilities, the alkalinization of the rhizosphere and the impact on auxin levels as potential sources of plant growth inhibition need to be further investigated.

## Acknowledgments

We thank Dr. Nishikant Wase of the W. M. Keck Biomedical Mass Spectrometry Laboratory at the University of Virginia School of Medicine for LC-MS quantification of AA in root exudates. This study was partially supported by funds from the University of Virginia through a 4-VA grant awarded to CHD.

## Author Contributions

IDKA and CHD conceived and designed the experiments. IDKA and CHD performed experiments with help from BTK. IDKA analyzed the data with input from CHD. IDKA and CHD wrote the manuscript. All authors read and approved the final version for publication.

## METHODS

### Plant materials

For all experiments, Arabidopsis wild-type plants were Col-0 and mutants were of Col-0 background. All T-DNA lines belong to the SALK Collection (60) and were obtained from the Arabidopsis Biological Resource Center, Ohio State University. Homozygous mutant lines were selected by PCR genotyping as previously described (Zhang et al 2022a). Seeds were always surface-sterilized using 10% bleach three times for two minutes, followed by three washes with sterile water, and resuspended in 0.1% PhytoAgar (PlantMedia; Cat#40100072-2). and stratified in the dark at 4 °C for at least two days.

### Growth of seedlings in vertical plates

Stratified seeds were germinated on square plates (100 mm × 100 mm square plates; Fisher Scientific; Cat#FB0875711A) containing sterile full-strength (1x) Murashige and Skoog (MS) basal medium with vitamins (PhytoTech # M519), supplemented with 0.5% sucrose (Sigma; S7903), 0.5 g/L MES (Sigma; M8250), and 0.7% PhytoAgar (PlantMedia #40100072-1). The pH was corrected to 5.7 with KOH. Plates were sealed with parafilm and incubated vertically in a reach-in plant growth incubator (Conviron Adaptis 1000, Canada) at 25 ± 0.2 °C, 75% RH, 16h Light/8h Dark, and 100 μmoles/m^2^/s light intensity for the times indicated in the relevant figure legends. Uniformly growing seedlings were then selected and transferred to new plates containing autoclaved-3MM paper cut to fit 100 mm x 100 mm square plates and wetted to saturation with 0.5x liquid MS medium pH 5.7 without sucrose. These plates were incubated horizontally under the same conditions as above for one day to allow roots to attach to the 3MM paper.

### Root exudate collection assays

Root exudates were collected using a modification of a previously published method(31). Arabidopsis seeds were sown on an autoclaved polytetrafluoroethylene (PTFE) mesh (McMaster-Carr; Cat#1100t41) floating on the surface of 1 mL 1x MS liquid medium containing 0.5% sucrose in 12-well tissue culture plates (USA Scientific; Cat#CC7682-7512). Plates were incubated in reach-in plant growth incubators (Conviron Adaptis 1000, Canada) at 25 ± 0.2 °C, 75% RH, 16h Light/8h Dark, and 100 μmoles/m^2^/s light intensity. Sterilized and stratified seeds were sown on the floating mesh. Twelve days after sowing, the medium was replaced with 0.5x MS liquid medium without sucrose, and plants were allowed to grow for three additional days. Importantly, in these conditions, roots grow submerged in medium while shoots remain on the air space of the well. Root exudates were then collected and filter-sterilized through 0.22 μm filter for further processing.

### Colorimetric/Fluorometric Quantitation of Amino Acids

Total amino acid content of root exudates was assessed using the L-Amino Acid Quantitation Colorimetric/Fluorometric Kit (BioVision #K639-100) following the manufacturer’s instructions with modifications. Briefly, a 12.5 μL reaction mix containing 11.5 μL L-amino acid assay buffer, 0.5 μL L-amino acid probe, and 0.5 μL L-amino acid enzyme mix was added to each well containing 12.5 μL of the test samples, L-amino acid standards to generate a standard curve, or unplanted media to estimate and later subtract the background signal. The reactions were incubated in the microtiter plate reader (SpectraMax^®^ i3x, Molecular Devices) for 30 minutes at 37 °C. The fluorescent signal (Ex/Em = 535/587 nm) was recorded every 5 min. AA quantification was performed on 5-6 independent samples per condition.

### Liquid Chromatography—Mass Spectrometry Analysis of Amino Acids

Root exudates were vacuum-dried and reconstituted in 100 μL of 0.1% formic acid. Samples were analyzed at the University of Virginia-Biomolecular Analysis Facility Core following a standard protocol (Nemkov et al., 2019). Serial dilutions of standards of every AA were ran for every analysis. Five μL of samples were injected into a UHPLC system (Ultimate 3000, Thermo, San Jose, CA, USA) and separated through a 3-min isocratic elution (5% acetonitrile, 95% water, 0.1% formic acid) on a 1.7 μm C18 column (Kinetex XB-C18, Phenomenex, Torrance, CA, USA) at 250 μL/min and 25°C. High-resolution mass spectrometry analysis was performed using a triple quadrupole orbitrap mass spectrometer (Q-Exactive HF-X, Thermo Scientific). Glycine data was omitted from final analysis as unplanted MS medium contained background levels of glycine (2 mg/L). Data analysis was performed with the proprietary Thermo XCalibur software (https://www.thermofisher.com/order/catalog/product/OPTON-30965#/OPTON-30965), and targeted peak detection was achieved using ICIS peak integration algorithm. Thermo Quantitative analysis software (Quan Browser) was then used to generate calibration curves, followed by determining the concentration of the amino acids in the root exudate and unplanted samples. LC-MS analysis was performed on 6 independent samples per condition.

### Bacterial growth on root exudates

*Pseudomonas simiae* WCS417r was maintained on LB plates supplemented with 50 μg mL^-1^ rifampicin. A single colony was randomly picked from a plate and grown overnight in 100 mL of LB at 28 °C and 230 rpm till the cultures reached OD_600_ = 0.4 – 0.8. The bacteria were then harvested by centrifugation, washed three times with sterile water, and the OD6_00nm_ was adjusted to the required inoculation titer with sterile water. Root exudates (100 μL) of each condition were aliquoted on six or more wells of a 96-well microtiter plate and inoculated with *Ps* WCS417r. Plates were incubated under constant agitation at 28 °C, and the culture density was monitored over time by reading OD_600nm_ in a microtiter plate reader (SpectraMax^®^ i3x, Molecular Devices).

### Root colonization

Seedings were grown on vertical plates as described above. Two weeks after sowing, six (6) uniformly growing seedlings were transferred to each of 5 plates containing 3MM paper wetted with 5 mL half-strength MS without sucrose. Plates were horizontally incubated for one day to allow seedlings to attach to the paper surface. The next day, the edge of the paper opposite to the seedling’s-shoot side was flooded with 0.5x MS alone (contamination control) or inoculated with freshly harvested *Ps* WCS417r diluted to OD_600nm_=0.0002. Bacteria were allowed to colonize the roots for 72h. The capacity of roots to support bacterial growth was analyzed in 6 to 30 individual seedlings per condition. To assess colony forming units (CFU) per mg of root, the root mass was recorded right after harvesting. Immediately after weighing, the root tissue was ground with metal beads in a TissueLyser (QIAGEN, Inc). Aliquots of the ground roots were base-ten serial diluted in sterile water and plated on OmniTray plates (Nunc^™^) containing LB agar medium supplemented with 50 μg/mL of rifampicin. Plates were incubated at 28 °C overnight, and colonies were counted under a microscope. Only replicates with no colonies in the “contamination control” were included in the analysis.

### Competitive chemotaxis assay

Competitive chemotaxis assay was designed by modifying capillary assays previously described (34, 38), using syringes (**Fig. S3**). Briefly, *Ps*WCS417r cultures grown and harvested as described above were resuspended at OD_600_=0.002 in chemotaxis buffer (10 mM potassium phosphate pH 7.2, 1 mM MgCl2, and 0.1 mM EDTA). Bacterial suspensions (40 mL) were pipetted into empty sterile Petri dishes. Then, 200 μL of the test samples (*e.g.,* unplanted MS, wild-type-root exudates, or *lht1*-root exudates) were loaded into each of two 1 mL sterile syringes (without needles) and the tips of the syringes were dipped below the liquid surface in the Petri dish containing the *Ps* WCS417r in chemotaxis buffer suspension. The syringes loaded with exudates were allowed to attract bacteria from the suspension for 30 min. The syringes were removed from the bacterial suspension, and colony-forming units in the exudates contained in the syringes were assessed via dilution and plating as described above (see root colonization assay for details).

### Biofilm formation assays

Unplanted medium, wild-type root exudates, and *lht1* root exudates were tested for their capacity to promote biofilm formation upon *Ps* WCS417r inoculation as follows. Each well of a 96-well round-bottom plate (Catalog # CLS2797, Sigma) was first loaded with 200 μl of root-exudate sample and then inoculated with 2 μL of freshly harvested *Ps* WCS417r (see details above) resuspended at OD_600_=0.02. Biofilms were allowed to grow statically for 48h at 28 °C. Subsequently, the medium and non-adherent bacterial cells were carefully removed by pipetting and wells were washed with sterile water without disturbing the biofilms attached to the bottom of the plate. The plates were air-dried for 5-10 minutes. Next, the wells were stained with 125 μL of a 0.1% (w/v) solution of crystal violet in water for 15 minutes. The stain was gently rinsed with water, without disturbing the stained biofilm. One hundred and fifty (150) μL of a 6:3:1 sterile water:methanol:acetic acid solution was added to each well to solubilize the crystal violet for 15 minutes. To quantify biofilm biomass, 125 μL of the 150 μL solution of stained biofilm were transferred into a well of a polyvinyl chloride 96-well flat-bottom plate (Catalog # CLS2595, Sigma), for absorbance assessment at 550 nm in a microplate reader (SpectraMax^®^ i3x (Molecular Devices). Biofilm formation was tested on 3 to 6 independent samples per condition.

### Plant growth on exudates ± *Ps* WCS417r

The capacity of *Ps* WCS417r to promote the growth of Arabidopsis seedlings when combined with WT exudates, *lht1* exudates, or WT exudates supplemented with Gln or Ser was assessed as follows. Seedings were grown on 1x MS PhytoAgar with 0.5% sucrose. Plates were incubated in reach-in plant growth incubators (Conviron Adaptis 1000, Canada) at 25 ± 0.2 °C, 75% RH, 16h Light/8h Dark, and 100 μmoles/m^2^/s light intensity. After two weeks, six (6) uniformly growing seedlings were transferred to each of 3 to 7 plates containing 3MM paper wetted with 5 mL half-strength MS without sucrose. Plates were horizontally incubated for one day to allow seedlings to stabilize. The next day, the seedlings in each plate were root flooded (liquid added to the edge of the paper opposite to the seedlings-shoot side) with half-strength MS alone (contamination control) or with freshly harvested *Ps* WCS417r diluted to OD_600nm_= 0.2 in the following carriers: Plate 1) WT exudate, and Plate 2) *lht-1* exudate. In independent experiments we tested half-strength MS alone as contamination control or *Ps* WCS417r diluted to OD_600nm_= 0.2 in the following carriers: Plate 1) WT exudate, Plate 2) WT exudate supplemented with 1mM Gln, Plate 3) WT exudate supplemented with 10mM Gln, Plate 4) WT exudate supplemented with 1mM Ser, and Plate 5) WT exudate supplemented with 10mM Ser. We also included an “overgrowth control” in which we root-flooded the seedlings with *Ps* WCS417r diluted in WT exudate at final OD_600nm_= 1.6 (instead of 0.2); this control was included to test whether overgrowth of *Ps* WCS417r would be sufficient to inhibit plant growth.

### *Ps* WCS417r excreted-ammonium assessment

The capacity of *Ps*WCS417r to produce ammonia through metabolizing exudates of interest was measured using a modified version of the Berthelot colorimetric method (**Fig. S4**). Briefly, 24 wells of a 96-well plate were filled with 90μl of WT or *lht1* exudate, or 0.5x MS media as control. Twelve wells of each condition and of each independent source of exudate (*e.g.,* 12 wells of WT exudate obtained in biological replicate 1, 12 wells of WT exudate obtained from biological replicate 2, etc.) were inoculated with freshly harvested and three times washed *Ps*WCS417r to reach a final OD_600nm_ = 0.5, while the remaining 12 wells of each condition and biological replicate were left bacteria-free to measure the basal levels of ammonia in each exudate (negative controls). Plates were incubated at 28°C with constant shaking. After 5, 10, 20, and 30 minutes, the content of 3 wells per condition was transferred to 1.5 mL tubes and centrifuged for 15 min at 15,000g. The 3 supernatants from each treatment and control condition were pooled and filtered sterilized. Two microliter aliquots of the supernatants were transferred to wells of a 96-well plate prefilled with 98μl of assay buffer of the ammonia assay kit from AbCam (AbCam Cat # ab102509), and ammonia levels were quantified according to the manufacturer’s guidelines. Each supernatant was tested in triplicate, and, as described above, three independent exudates per condition were tested in parallel.

### Plant tissue gene expression analysis

Root/leaf tissues were harvested one at a time into previously weighed tubes, and the weight of the tissues was measured. Immediately after weighing a sample, it was flash-frozen in liquid nitrogen and kept at −80°C until processing. RNA was isolated using TRIzol^®^ Reagent (Fisher Scientific; Cat#15596018), DNAse-I treated (Promega) and quantified in a NanoDrop-ND1000 spectrophotometer (Thermo Fisher Scientific Inc). Two (2) μg of total RNA were used for cDNA synthesis with MMLV (Promega) and random hexamers (Invitrogen). qPCR was performed using real-time SYBR-Green quantification. There were two technical replicates per condition per biological replicate. The reactions were performed an ABI 7500 Fast Real-Time PCR system (Applied Biosystems). Data were analyzed using the ΔΔCt method (61). Gene expression was measured on ≥3 independent samples per condition.

### Statistical analysis

Data analyses and graphs were generated using Excel and GraphPad. A two-sided Student’s *t*-test was performed for statistical comparison of two means, or a Welch’s *t*-test for two means with unequal variances, when relevant. For comparison of more than two means, a one-way ANOVA followed by Tukey’s posthoc test, or a Kruskal-Wallis test for unequal variances followed by Dunn’s posthoc test was performed, as indicated in the relevant figure legends.

For statistical analysis of bacterial growth curves, the CGGC (Comparison of Groups of Growth Curves) permutation test (62) was used to compare pairs of samples (i.e., unplanted vs. wild type; unplanted vs. *lht1;* wild type vs. *lht1)* over the course of growth (24 hours). The test statistic (mean *t*) is the two-sample *t*-statistic to compare the OD_600_ values between the two groups at each hour, averaged over the course of growth (24 hours). A P-value was obtained for the test statistic by simulation. Samples were randomly allocated to each of the two groups and the mean *t* was recalculated for 10,000 data sets generated through this permutation. The P-value is the proportion of permutations where the mean *t* is greater in absolute value than the mean *t* for the original data set.

---

**Figure S1.**
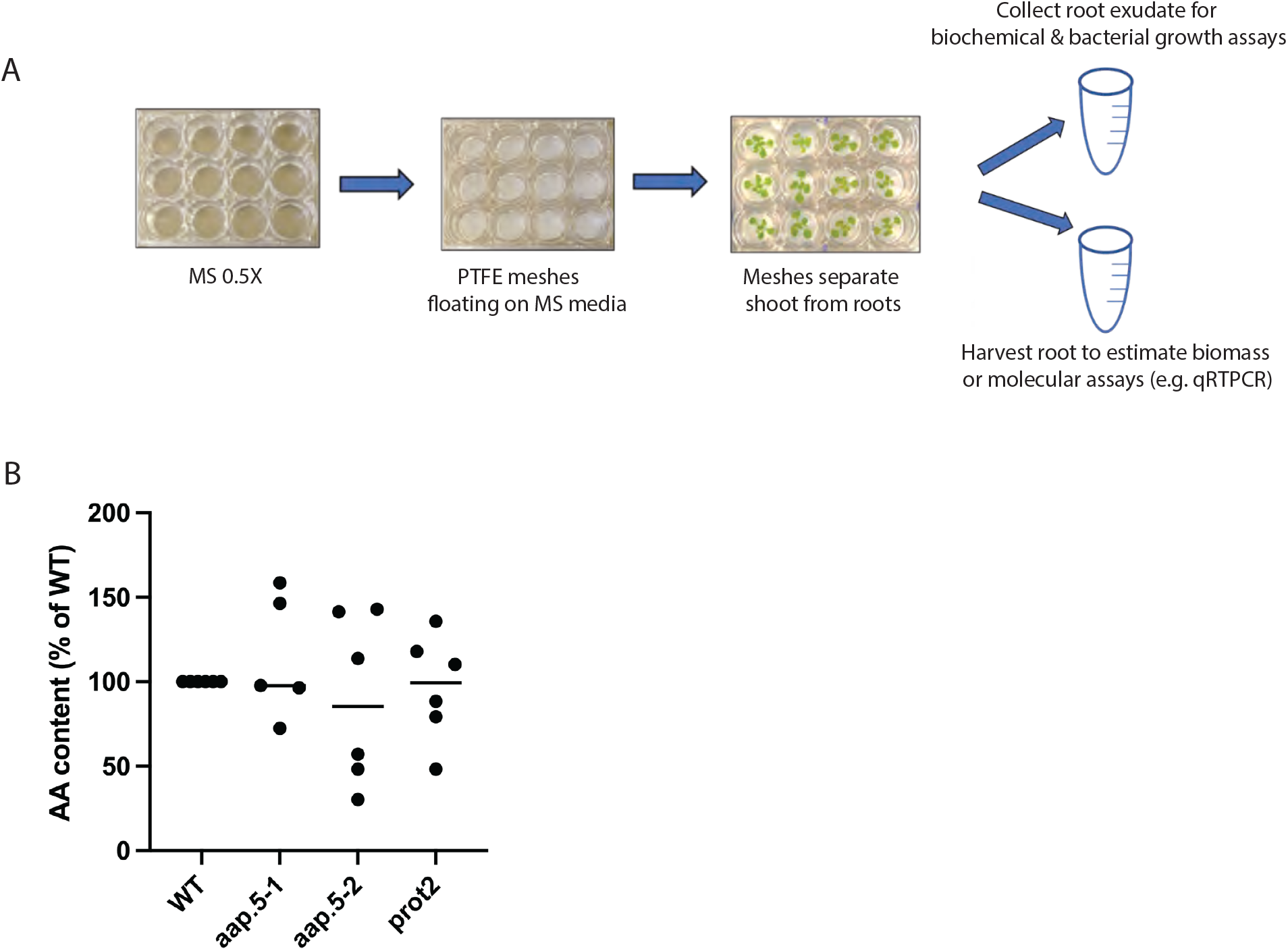
**(A)** Workflow to obtain root exudates. Twelve-well plates were loaded with 1 mL of 1x MS buffer supplemented with 0.5% sucrose and 0.5 g/L MES. Seeds were sown on sterile polytetrafluoroethylene (PTFE) meshes and placed floating on top of the medium. After 12 days, the medium was replaced with 0.5x MS liquid medium without sucrose, and plants were allowed to grow for three additional days. Importantly, in this condition, roots grow submerged in the medium while shoots remain in the air space of the well. Root exudates were then collected and filter-sterilized for further processing or testing. **(B)** Total AA content in *aap5* and *prot2* root exudates is similar to that of WT plants. Seeds of sequence-verified homozygous mutants were sown, and exudates were obtained as described in (A). Colorimetric quantification of AAs was carried out as described in Methods. Each dot corresponds to a biological replicate and is the median AA content per mg of roots of 5 seedlings per genotype relative to the wild type (as a percentage). Experiments were repeated 6 independent times. One-way ANOVA followed by Tukey’s posthoc test shows no statistically significant differences between WT and the mutant genotypes.

**Figure S2.**
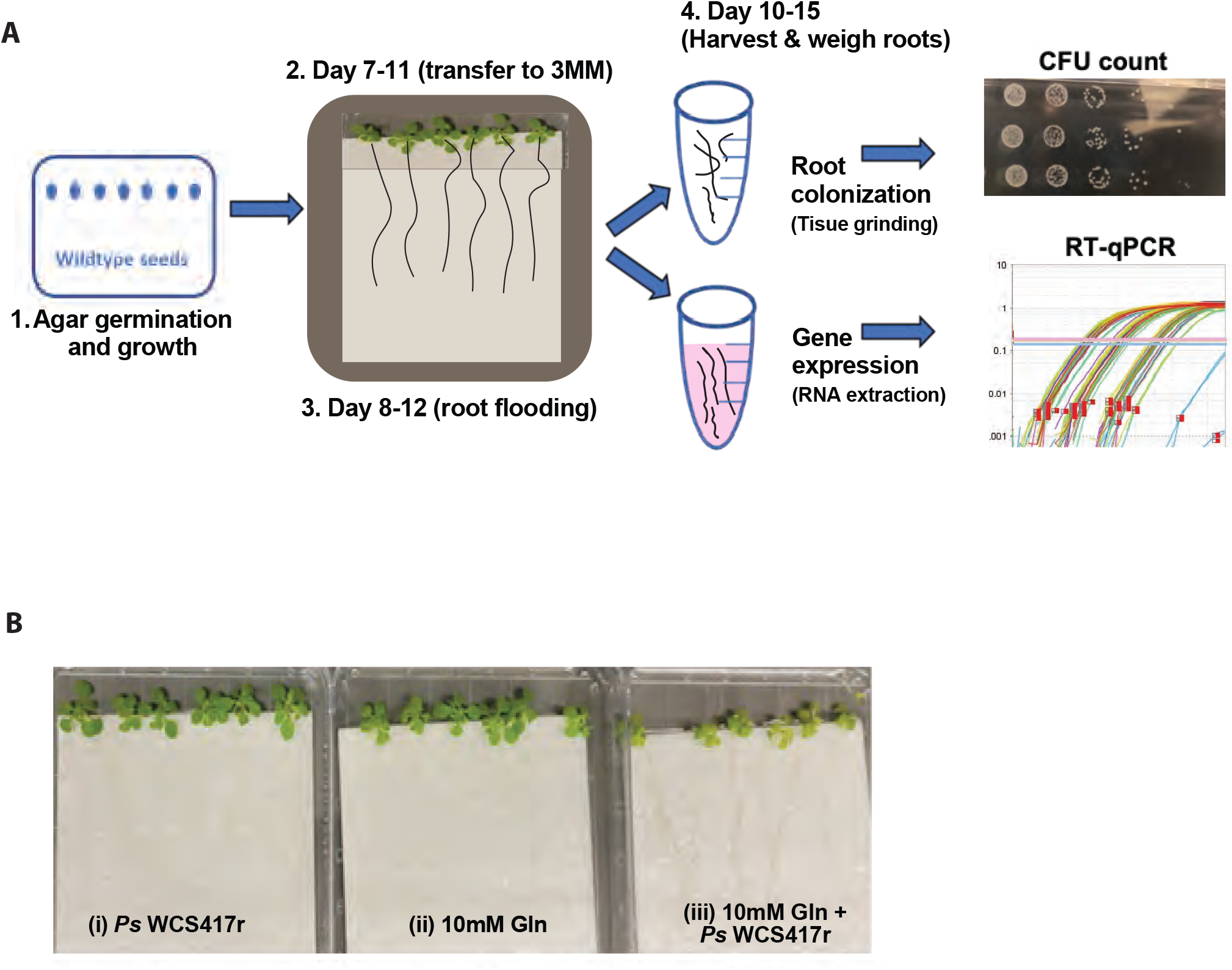
**(A)** Workflow to assess colonization and gene expression in roots. **(1)** Stratified seeds are germinated in 1x MS agar with 0.5% sucrose. **(2)** At days 7 to 11, uniformly growing seedlings were transferred to square plates containing autoclaved 3MM paper wetted with 0.5x MS without sucrose (composed image to reveal roots position and size). Plates were sealed and incubated horizontally for 1 day to allow roots to attach to the 3MM paper. **(3)** At days 8 and 12, the bottom of the 3MM paper is flooded with a suspension of *Ps* WCS417r (final OD600nm= 0.2) or sterile 0.5x MS as a no-growth (contamination) control. The bacteria are resuspended in exudates obtained from WT or lht1, and WT exudates were supplemented with AAs as indicated. 4. After 24-72h of exposure to exudates *Ps* WCS471r AAs, roots are harvested, weighed, and processed to obtain RNA or to count bacterial colony-forming units (CFU). (B) Representative pictures of seedlings in square plates on 3MM paper substrate. At day 13 roots were flood-treated with: (i) Ps WCS417r OD600nm=0.2 (control), (ii) 10 mM Gln, or (iii) *Ps* WCS417r and Gln at final OD600nm=0.2. The images show symptoms 3 days after treatment.

**Figure S3.**
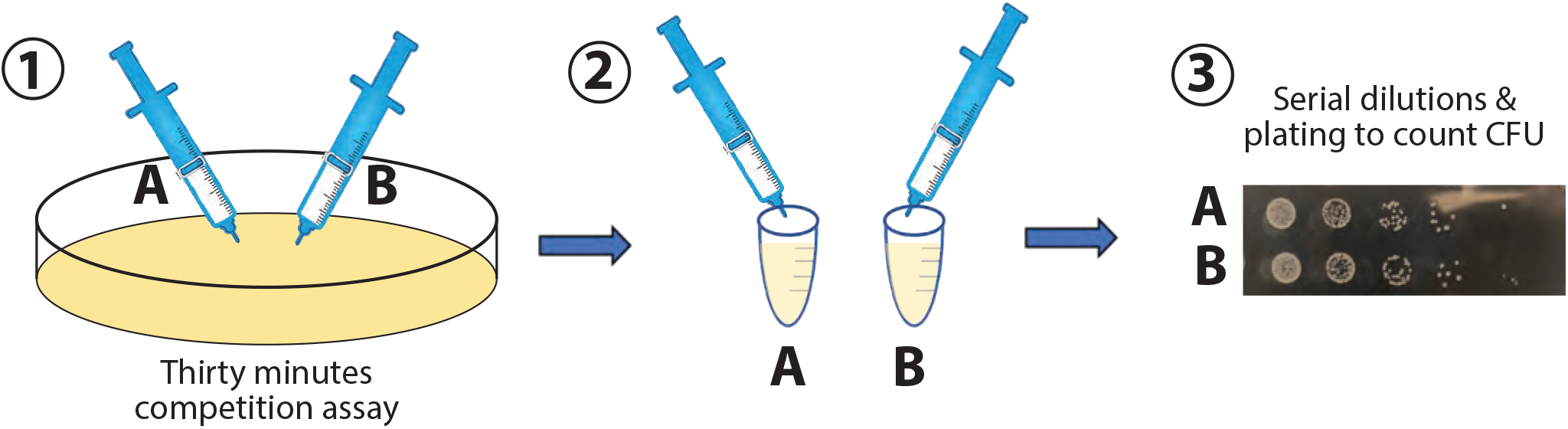
Workflow of competitive chemotaxis assay. **(1)** A Petri dish is filled with 40 mL of *Ps* WCS417r freshly resuspended chemotaxis buffer at a final OD600nm =0.002 Filter-sterilized root exudates (200 μL) or fresh MS medium (negative control) were loaded into 1 mL sterile syringes (without the needles). The tips of the two syringes to be tested in the competitive assay were then immersed just under the surface of the bacterial suspension contained in the Petri dish. **(2)** After 30 min, the content of each syringe was transferred to independent 1.5 mL tubes. **(3)** Serial dilutions were plated on LB-agar plates to count colony-forming units (CFU).

**Figure S4.**
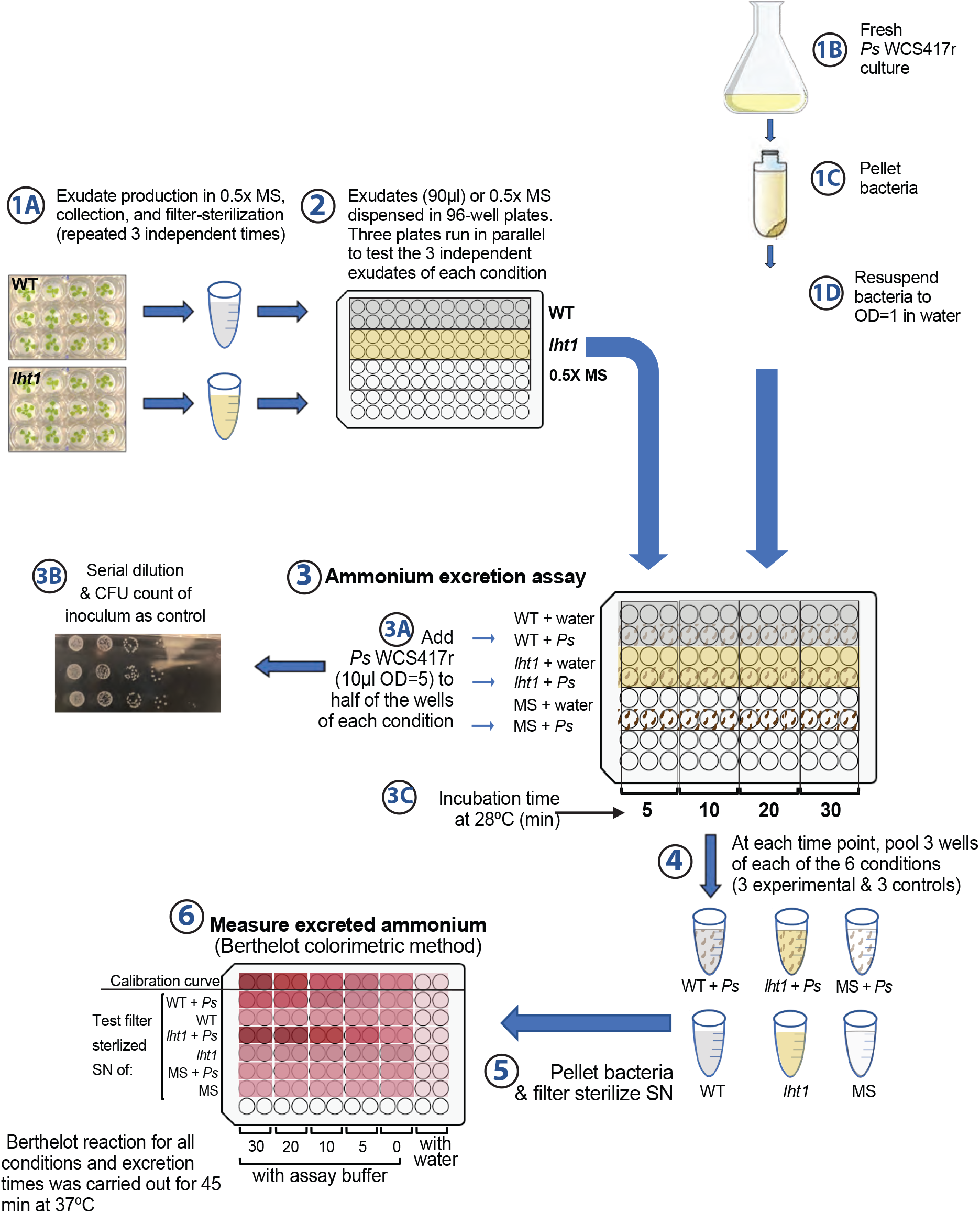
Workflow of *Ps* WCS417r excreted-ammonium assay. WT and *lht1* exudates were collected **(1A)** as described in Methods. Aliquots of 90 μL of each exudate were transferred to 24 wells of a 96-well plate **(2)**. While 12 wells of each exudate were inoculated with freshly harvested **(1A)** and three times washed **(1B)** *Ps* WCS417r to reach a final OD600nm = 0.02 **(1C)**, the remaining 12 wells of each condition were left bacteria-free to be used as negative controls. Plates were incubated at 28 °C with constant shaking **(3)**. At 5, 10, 20, and 30 minutes, 3 wells of each experimental and 3 wells of each control condition were harvested **(3A)**. In parallel, 10 μL aliquots were used to assess CFUs (3B). After centrifugation for 5 min at 10,000g, the 3 supernatants from each treatment and control condition were pooled **(4)** and filtered-sterilized **(5)**. Aliquots of the supernatants were used to quantify ammonia **(6)** using the ammonia assay kit from AbCam (Cat # ab102509), according to the manufacturer’s guidelines. The experiment was repeated 3 times with independent batches of fresh exudates.

